# Annelid perspectives into the role of FGF signaling in caudal regeneration

**DOI:** 10.1101/2024.11.26.625069

**Authors:** Alexandra Y. Shalaeva, Vitaly V. Kozin

**Affiliations:** Department of Embryology, St. Petersburg State University, St. Petersburg, Russia

**Keywords:** regeneration, FGF, annelids, blastema, mesoderm, caudal, even-skipped, polarity, AP axis, growth zone, segmentation

## Abstract

Fibroblast growth factor signaling plays a crucial role in various developmental processes and is a key driver of regeneration. In annelids this pathway is active from the earliest stages of reparative morphogenesis, yet its specific function remains unclear. Here we have functionally examined FGF signaling following the amputation of posterior segments in the marine annelid *Alitta virens*. We utilized the pharmacological agent SU5402 to inhibit the FGF receptor kinase at different time points. With whole-mount in situ hybridization we analyzed the expression of regulatory genes that pattern posterior territories (*cdx, evx, post2*), multipotent/germ cells (*vasa, piwi*), mesodermal tissues (*twist*), and segmental boundaries (*engrailed*). Our findings reveal that FGF signaling is essential for blastema induction by promoting dedifferentiation and proliferation of cells at the wound site but is not involved in posteriorization. On the contrary, this pathway is crucial for differentiation in the anterior part of the regenerative bud, impacting its mesodermal derivatives and segment boundary formation. Comparative analysis suggests that while certain functions of FGF signaling, particularly in mesodermal patterning, are conserved across taxa, its role in posterior axis elongation appears to have evolved specifically within the vertebrate lineage. This research enhances our understanding of the evolutionary origins and functional diversification of FGF signaling in regeneration, positioning *A. virens* as a valuable model for exploring the complexities of regenerative biology.

## Introduction

Post-traumatic regeneration represents a complex and fascinating process that gained an increased attention in recent decades (Bideau et al., 2021; Lai & Aboobaker, 2018; Mehta & Singh, 2019; Rennolds & Bely, 2023; Srivastava, 2021). Regeneration abilities at the organismal level vary even between closely related phylogenetic lineages demonstrating significant evolutionary plasticity (Elchaninov et al., 2021; Zattara et al., 2019; Zattara & Bely, 2016). Some animals are incapable of regeneration at all, some can only partially restore certain cell types or tissues, while others can regrow their entire body from just a small fragment (Bely & Nyberg, 2010). These fundamental differences are associated with the regulation of specific mechanisms and interdependent steps of regeneration (Liu et al., 2021; Tiozzo & Copley, 2015). The most well-established whole-body regeneration models include cnidarians and planarians, which possess unique stem cell populations (Reddien, 2018; Reddy et al., 2019; Röttinger, 2021). Annelids on the other hand have a much more complex body plan and tissue organization, making them an interesting model for Evo-Devo regeneration studies (Özpolat & Bely, 2016; Nikanorova et al., 2020; Kostyuchenko & Kozin, 2021; Zattara & Rennolds, 2024; Zattara, 2020; Özpolat, 2023). Regeneration in the majority of aforementioned organisms happens in the similar sequence of stages: wound healing, formation of the wound epithelium, which induces the blastema, followed by blastema growth, patterning, morphogenesis and differentiation of its cells. And for these stages regardless of the organism fundamental questions are cellular sources and molecular drivers of the process (Bideau et al., 2021; Galliot, 2014; Guerin et al., 2021; Tanaka & Reddien, 2011).

Fibroblast growth factors (FGFs) are ancient signaling molecules that have a wide range of actions, and involved in diverse processes of development and regeneration (Bertrand et al., 2014; Farooq et al., 2021; Maddaluno et al., 2017; Ornitz & Itoh, 2015, 2022). This diversity of functions is accomplished by an extensive repertoire of ligands and receptors, which varies among different organisms, reaching up to 22 ligands, grouped into 7 subfamilies, and 4 receptors (FGFR) in vertebrates (Ornitz & Itoh, 2022). Moreover, FGFR’s variable specificity for binding ligands can be amplified by alternative splicing (Rebscher et al., 2009; Zhang et al., 2006). FGFR’s intracellular targets include the mitogen-activated protein kinase/extracellular signal-regulated kinase pathway (MAPK/ERK), as well as phosphatidylinositol 3-kinase/protein kinase B (PI3K/AKT), and phospholipase C gamma (PLCγ) (Böttcher & Niehrs, 2005; Eswarakumar et al., 2005; Ornitz & Itoh, 2015). Such a wide range of downstream pathways for signal transduction provides FGFs with numerous mechanisms for regulating cellular behavior, e.g., promoting proliferation, apoptosis, migration, and differentiation (Dailey et al., 2005; Dorey & Amaya, 2010; Ornitz & Itoh, 2015; Tulin & Stathopoulos, 2010). Naturally, the FGF pathway operates in major developmental processes in metazoans, such as cell fate specification, gastrulation, somitogenesis, axial elongation, and morphogenesis (Böttcher & Niehrs, 2005; Thisse & Thisse, 2005; Tulin & Stathopoulos, 2010; Dorey & Amaya, 2010; Green et al., 2013; Fan et al., 2018; Andrikou & Hejnol, 2021; Kumar et al., 2021).

The most well-known role of FGF signaling in mesoderm induction. It was first demonstrated in the *Xenopus* embryo, where basic FGF (FGF2) was proposed to function as a morphogen in mesoderm formation (Kimelman & Kirschner, 1987; Slack et al., 1987), which was confirmed by further genetic manipulations (Amaya et al., 1991). A similar function was later described in other vertebrates (Dorey & Amaya, 2010; Kumar et al., 2021) and in invertebrates (Andrikou & Hejnol, 2021; Chou et al., 2024; Fan et al., 2018; Seudre et al., 2022; Tulin & Stathopoulos, 2010). Another prominent role of FGF signaling is in axial patterning, affecting specifically the posterior elongation of the neural tube and mesodermal derivatives in vertebrate embryos (Boulet & Capecchi, 2012; Griffin et al., 1995). The confined expression of fgf8 in the embryo’s posterior tip maintains neuro-mesodermal progenitor cells in an immature state (Diez del Corral & Morales, 2017). The anteroposterior (AP) gradient of fgf8 activity, formed by mRNA decay, serves as both a posteriorizing factor and an instruction for the future differentiation of tissues (Chandel et al., 2023; Dubrulle & Pourquié, 2004b; Oginuma et al., 2017). The FGF molecules often act alongside other signaling pathways such as BMP and Wnt, together providing an opposing gradient to anteriorly produced retinoic acid, and thus establishing anterior-posterior (AP) positional information (del Corral & Storey, 2004; Dubrulle & Pourquié, 2004a; Wahl et al., 2007).

The annelid *Nereis diversicolor* was the first invertebrate in which fibroblast growth factors were identified, and their role in regeneration and growth was proposed (Blanckaert et al., 1992). Today annelids are emerging as a key model for regeneration research, owing to their complex body plan and, in the case of nereidid polychaetes, their robust and rapid reparative processes (Kozin et al., 2017; Metzger & Özpolat, 2024; Planques et al., 2019). This rapid regeneration is facilitated by active cellular proliferation, which begins after blastema formation. Blastemal cells in annelids arise either through dedifferentiation of cells in the tissue adjacent to the wound, as observed in nereidids (Bideau et al., 2024; Boilly, 1969; Herlant-Meewis & Nokin, 1962; Kozin & Kostyuchenko, 2015; Planques et al., 2019; Shalaeva & Kozin, 2023; Stockinger et al., 2024), or through the migration of specialized cells from other regions (de Jong & Seaver, 2018; Myohara, 2012; Zattara et al., 2016). Regardless of their origin undifferentiated (presumably multipotent) cells at the wound site proliferate extensively and restore the missing body parts. In previous work, we discovered that *Alitta virens* utilizes the FGF signaling pathway to promote cell cycle entry during posterior regeneration (Shalaeva et al., 2021). Both FGF ligands and receptors show an early response to amputation, being detected at the wound site as soon as 4 hours post-amputation (hpa). Following the formation of the regenerative bud, transcripts of both FGF ligands and receptors are present throughout the regenerating tissues. The *Avi-fgf8/17/18* and *Avi-fgfA* transcripts are found in ectoderm and mesoderm, while *Avi-fgfr1* is specifically expressed in blastemal cells, and *Avi-fgfr2* is expressed predominantly in the ectoderm of the bud and transiently in the early blastema. Immediate inhibition of FGFR after amputation completely prevents bud formation, while inhibition during blastema formation results in a bud that lacks oblique muscles, although the terminal region of the regenerate and pygidial muscle ring still differentiate (Shalaeva et al., 2021). These findings suggest that FGF signaling plays an evolutionarily conserved role in blastema induction in *A. virens*.

FGF molecules have been shown to induce blastema formation in a variety of animals (Farooq et al., 2021; Maddaluno et al., 2017; Nacu et al., 2016; Satoh et al., 2016; Shibata et al., 2016). Functional studies confirm this role primarily in vertebrates. In zebrafish fin regeneration, FGF from the wound epithelium stimulates FGFR-expressing mesenchymal cells to form a blastema (Shibata et al., 2016). In the fiddler crab *Uca pugilator*, FGF2, which originates from the severed nerve, regulates mitotic activity in the blastema, driving its growth (Hopkins, 2001). In vertebrate models, FGF-soaked beads can rescue regeneration in denervated amphibian limbs and tails (Makanae et al., 2016; Mullen et al., 1996), and nerve-dependent FGF release has also been observed in chick and mouse regeneration (Maddaluno et al., 2017; Satoh et al., 2018). In the planarian *Dugesia japonica* (Auwal et al., 2020; Cebrià et al., 2002; Ogawa et al., 2002), *Djfgf* regulates neoblast proliferation via the ERK pathway, acting in a proliferative post-blastema that supplies neoblasts for the differentiation of lost body parts (Auwal et al., 2020). Given the diversity of tissues utilizing FGF signaling during regeneration, an important question remains: What is the evolutionary origin of FGF functions? This question has not been rigorously tested in organisms with a complex body plan and high cell fate plasticity.

In the present study, we aim to elucidate the precise developmental roles of FGF signaling during *A. virens* regeneration, particularly in the context of molecular patterning. We treated experimental specimens with the pharmacological inhibitor SU5402 to block FGF receptor activity at key stages of regeneration, including wound closure, blastema formation, patterning, and segmentation (Table 1). We used whole-mount in situ hybridization (WMISH) method to analyze the expression patterns of seven genes associated with specific regulatory states and tissues in regenerating nereidid worms. Our data provide new insights into the conserved functions of the FGF pathway, such as blastema induction, regulation of mesodermal derivatives, and restoration of positional information along the AP axis.

**Table 1.**
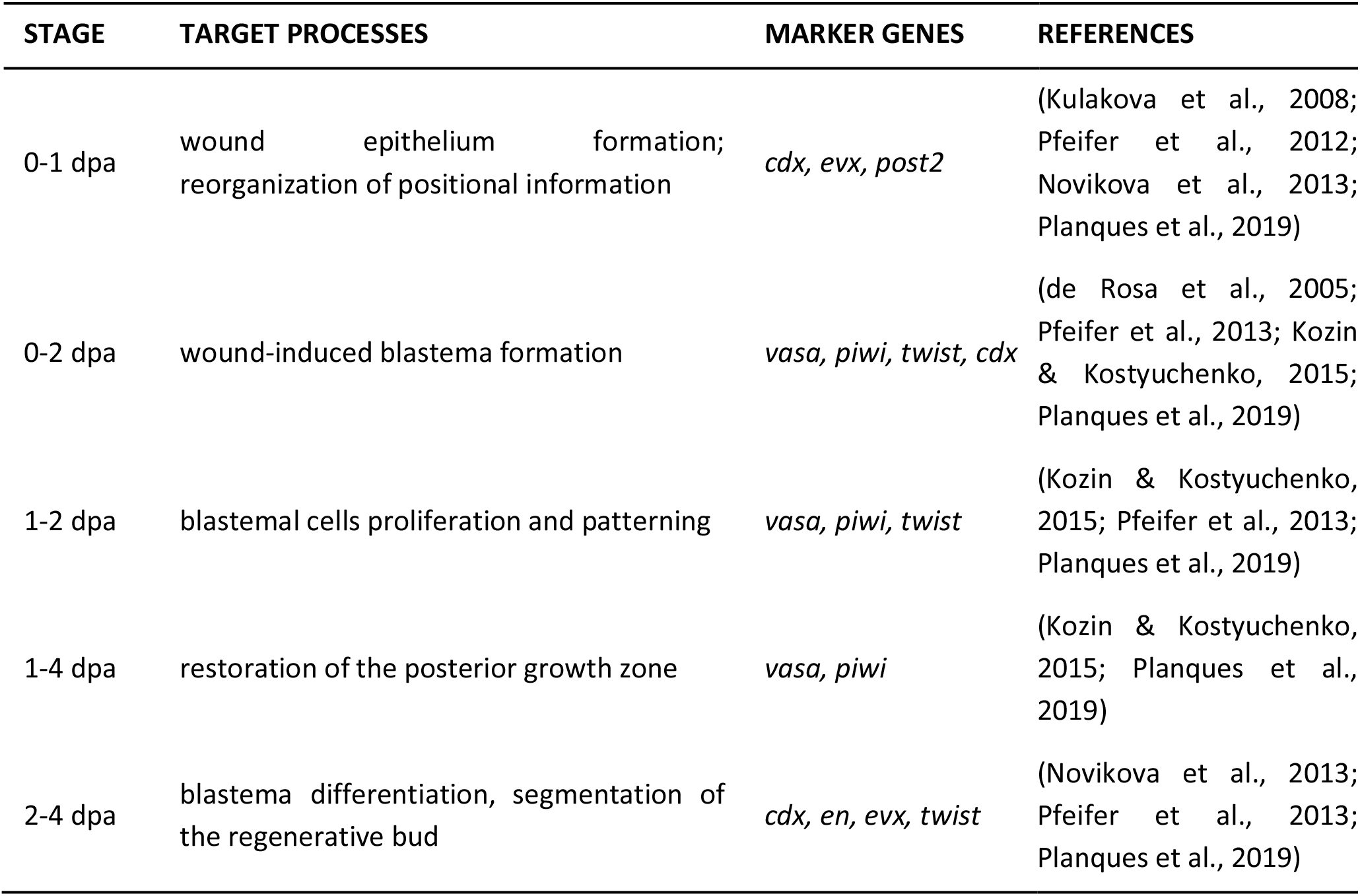
Overview of regenerative processes in nereidids after posterior amputation that were analyzed in the present work. We studied expression of seven marker genes that are associated with the listed processes. Suppression of the FGF pathway with SU5402 (inhibitor of FGFR tyrosine-kinase) was performed at the indicated stages.

## Materials and methods

### Animals

Spawning epitoke *Alitta virens* were caught in summer near the White Sea Marine station of St. Petersburg State University. A laboratory culture of embryos was obtained by artificial fertilization (Dondua, 1975). The animals grew for 2–3 months in small aquariums with natural or artificial seawater until they reached 15–20 segments in length. The posterior thirds of the juveniles’ bodies were amputated, and then the animals were left to regenerate for various time periods at +18 °C. At the preferred stages (1–4 days post-amputation, dpa), the specimens were anesthetized with 7.5% MgCl2 mixed with artificial seawater (1:1) and fixed in 4% paraformaldehyde on 1.75× PBS with 0.1% Tween-20 overnight at +4 °C. The samples were washed two times and stored in 100% methanol at −20 °C.

### SU5402 treatment

Pharmacological inhibition of FGF signaling by 50 μM of SU5402 (Sigma-Aldrich) in regenerating was standardized previously (Shalaeva et al., 2021). The powder was diluted with DMSO (producing stock concentration of 25 mM), aliquoted, and stored at −20 °C. In experimental setting SU5402 was mixed with artificial seawater, and then added to regenerating worms. In the case of prolonged incubation, the solution was changed every day.

### Whole-mount in situ hybridization

The samples were rehydrated from methanol, rinsed in PTW, treated with proteinase K (100 μg/mL) for 2.5–3 min at +22 °C, twice rinsed in glycine (2 mg/mL). Then samples were treated with 0,1M triethanolamine (TEA) pH = 8.0, and with solution of 0,1M TEA and acetic anhydride. Samples were postfixed with 4% PFA on PTW and washed in PTW before pre-hybridization. After overnight incubation with the probe and subsequent washes, the samples were blocked in 5% sheep serum on maleic acid buffer (MAB), followed by anti-digoxigenin AP antibodies overnight incubation (dilution 1:2000 in MAB). The staining was performed with NBT/BCIP, followed by washing in PTW and mounting in 90% glycerol. In situ hybridization results were visualized using DIC optics with Axio Imager D1 microscope (Carl Zeiss). Figure plates and schematic illustration were arranged in Inkscape.

## Results

### FGF pathway does not affect posteriorization

At the first day post amputation (dpa), molecular reestablishment of the new posterior end, i.e. posteriorization takes pace. This process reflects molecular morphallaxis, meaning that the molecular profile of the cells changes in response to their new relative position along the body axis (Kostyuchenko & Kozin, 2020). We analyzed genes that are involved in the posterior patterning, such as homeobox-containing *cdx/caudal, evx/even-skipped*, and *post2* (de Rosa et al., 2005; Novikova et al., 2013). mRNA of these genes was observed as early as 4 hpa at the wound site, which makes them one of the earliest activated genes in regenerative response (de Rosa et al., 2005; Novikova et al., 2013). FGFs are also activated by 4 hpa in *A. virens* (Shalaeva et al., 2021). In the 1 dpa DMSO controls, *Avi-cdx* was expressed in the wound epithelium and in the wound-adjacent gut (Fig. 1, arrows). A similar pattern was found for *Avi-evx*, however it was also weakly expressed in several groups of the ventral nerve cord (VNC) cells (out of focus in the control specimen in Fig. 1). *Avi-post2* mRNA was identified along the entire VNC ganglion with gradual increase in signal towards the wound site (Fig. 1, arrowheads). Apart from VNC, *Avi-post2* was expressed in sub-ectodermal bilateral domains under the wound epithelium (Fig. 1, arrows), and in parapodia. After inhibiting FGFR we recovered all these expression domains, without visible disruption of their localization at 1 dpa (Fig. 1 lower row). Thus, the earliest morphallactic events and wound healing in *A. virens* are independent of FGF signals.

**Figure 1.**
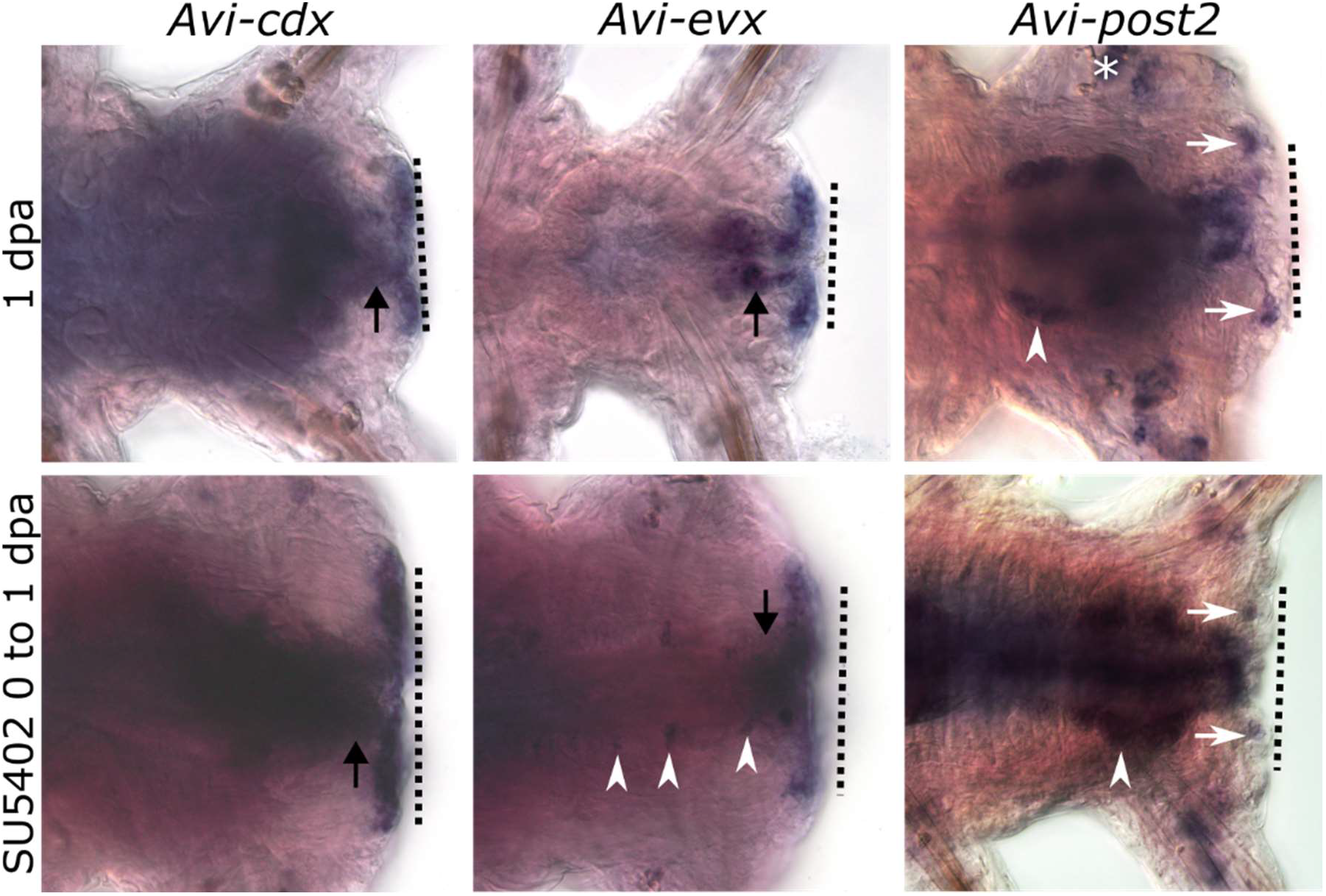
Posterior regeneration at 1 dpa stage after inhibition of FGF pathway (lower row), gene expression remains unchanged in comparison with DMSO controls (upper row). All planes are ventral views. Dotted line indicates amputation plane. In *Avi-cdx* and *Avi-evx* arrow indicates signal in the gut, arrowheads in *Avi-evx* and *Avi-post2* mark expression in the VNC, arrow in *Avi-post2* points to the sub-epithelial terminal domain. Asterisks mark non-specific staining.

### FGF pathway is necessary for induction of blastema and gain of mesodermal identity

The regeneration blastema forms in *A. virens* by 2 dpa stage (Kozin et al., 2017; Shalaeva & Kozin, 2023). The putative source of blastema is associated with de novo expression of dedifferentiation marker genes, such as *vasa* and *piwi*, which encode RNA-binding proteins. These molecules establish an evolutionary conserved germline multipotency program (GMP), characteristic for multipotent undifferentiated cells (Juliano et al., 2010; Kozin & Kostyuchenko, 2015). A bHLH TF-encoding gene *twist* is a blastemal marker, which is expressed later in regeneration process and is specific for mesodermal derivatives (Barnes & Firulli, 2009; Kozin et al., 2016; Pfeifer et al., 2013). We also analyzed expression of the aforementioned *Avi-cdx* as a marker for pygidial ectoderm (Kulakova et al., 2008). The most prominent effect of SU5402 on gene expression at 2 dpa was observed for *Avi-twist* and *Avi-piwi. Avi-twist* is normally found in the muscle-forming mesodermal part of the regenerative bud (Fig. 2). After exposition in SU5402 at 0–2 dpa *Avi-twist* expression was completely absent that was expected due to the lack of epimorphic regenerative bud (this phenotype was previously described in Shalaeva et al., 2021). But if we postpone the start of inhibition by one day after the amputation (exposition in SU5402 at 1–2 dpa), *Avi-twist* mesodermal domain was recovered in the miniature bud (Fig. 2, arrows). We observed similar effect for *Avi-piwi*. Normally, at 2 dpa stage, this gene is activated in the epidermal and mesodermal parts of the regenerative bud (Fig. 2). After inhibition at 0–2 dpa the gene expression was not detected, and similarly to *Avi-twist* the first day without FGFR suppression is enough to activate *Avi-piwi* at least partially (Fig. 2). In this experimental design we noticed *Avi-piwi* mRNA signal in the longitudinal muscle cell layer abutting the wound (Fig. 2, arrowhead), which is not found in the control specimens.

**Figure 2.**
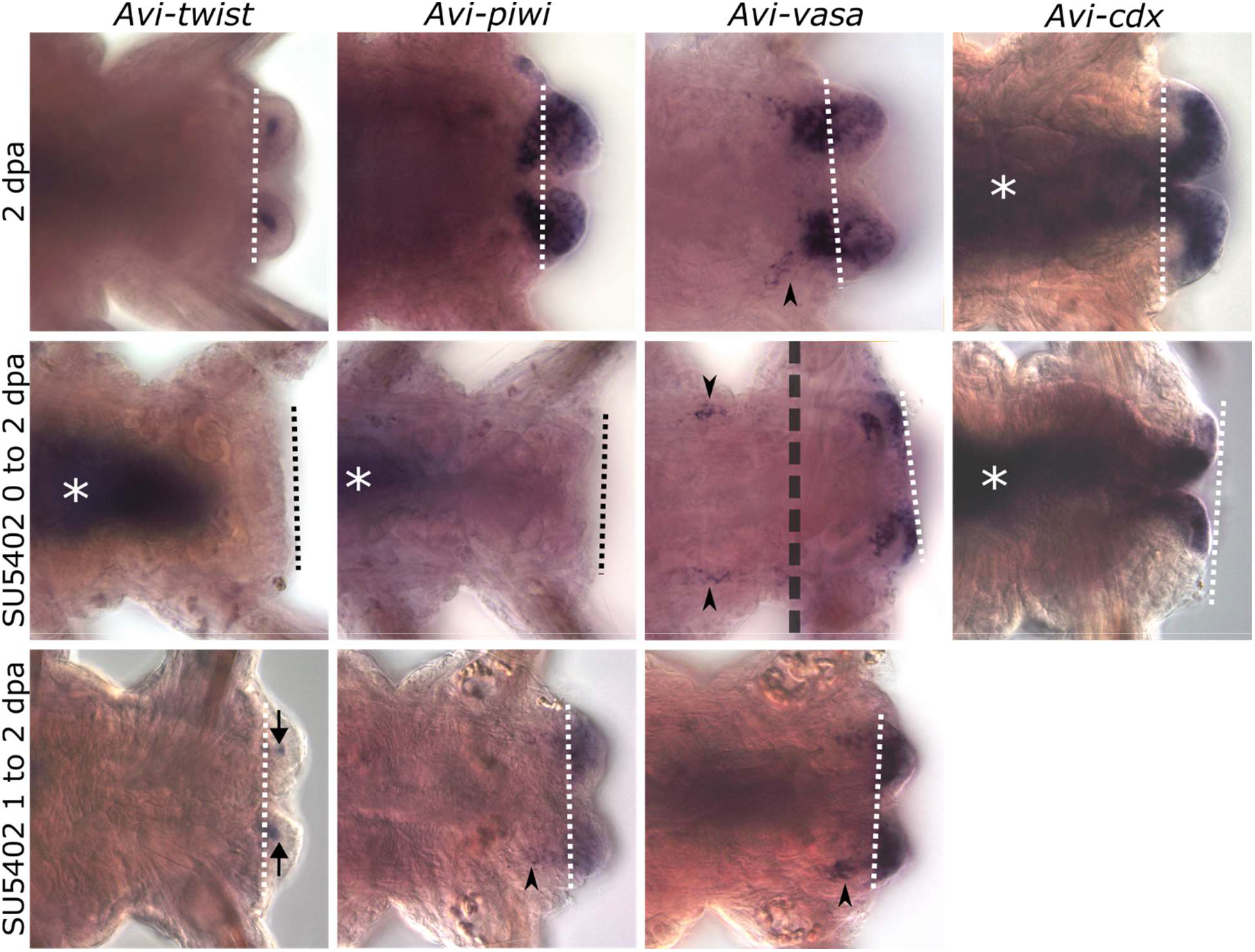
Posterior regeneration at 2 dpa stage in DMSO controls (upper row) and after inhibition of FGF pathway starting immediately after amputation (middle row) or at 1 dpa stage (lower row). All planes are ventral views. Dotted line indicates amputation plane. Arrows indicate reduced *Avi-twist* expression in the blastema, arrowheads in *Avi-piwi* and *Avi-vasa* mark expression within the muscle cell layer. In *Avi-vasa* in the middle row two optical planes of the same specimen are divided by grey dotted line. Asterisks mark non-specific staining.

In contrast to *Avi-twist* and *Avi-piwi, Avi-vasa* activated its expression in the wounded segment irrespective of SU5402 treatment. *Avi-vasa* domains in control specimens were found in epidermis and in blastemal cells, as well as in the muscles abutting the wound (Fig. 2, arrowhead). This pattern was completely reproduced in the reduced regenerative bud after inhibition at 1–2 dpa. Unexpectedly, after exposition in SU5402 at 0–2 dpa, activation of *Avi-vasa* was observed in the wounded segment, and even to a greater extent than is observed normally at 1 dpa (Kozin & Kostyuchenko, 2015). *Avi-vasa* expression was expanded in the epithelium and in internal cells below it (Fig. 2), which correspond to blastemal precursors within the muscle layer. In the same manner, for *Avi-cdx* after exposition in SU5402 at 0–2 dpa, an advanced pattern in comparison to 1 dpa, but not identical to the control was observed at 2 dpa. The strongest *Avi-cdx* expression marked the posterior parts of the gut and the abutting epithelium. Noteworthy, *Avi-cdx* in experimental specimens was downregulated in the superficial tissues, which produce the pygidial region in controls, at shorter distance from the gut opening (Fig. 2).

Summarizing the experimental results, we noticed that the tested regulatory genes had different responses to the FGFR inhibition in long-term (0–2 dpa) vs. short-term (1–2 dpa) exposure. When using SU5402 immediately after amputation, in contrast with normal expression, we observed either the absence of expression (*Avi-twist* and *Avi-piwi*), or the lack of drastic changes (*Avi-vasa* and *Avi-cdx*). Intriguing differences in the response of the two GMP markers *vasa* and *piwi* suggest a multi-stage and complex regulation of the dedifferentiation process. The short-term (1–2 dpa) exposure uniformly results in minimal effect on the miniaturized expression domain that indicates the utmost importance of the first 24 hours after injury for the induction of regeneration cell sources.

### FGF pathway in blastema differentiation and restoration of the posterior growth zone

Other set of experiments addressed the role of FGF pathway in differentiation of blastema, its growth and segmentation. Following SU5402 treatment from 2 to 4 dpa we tested expression of *Avi-twist* (mesodermal marker), *engrailed/Avi-en* (marker of the segmental boundaries (Kairov & Kozin, 2023)), *Avi-evx* and *Avi-cdx* (posterior markers). For *Avi-twist* and *Avi-en* we found significant pattern disturbances, whereas *Avi-evx* and *Avi-cdx* were expressed after inhibition like in controls.

*Avi-twist* expression pattern once again demonstrated the most prominent response to experimental manipulation. Normally, at the 4 dpa stage, its domains are found in the mesoderm. One of them is located where new segments are formed, and has a metameric character, that becomes more prominent with the growth of the bud. Another *Avi-twist* domain is found in the forming circular pygidial muscles of the anal sphincter (Fig. 3A, arrowhead). When we inhibited FGFR at 2–4 dpa interfering with blastema differentiation, the posterior-most pygidial mesodermal domain was the only one remaining (Fig. 3A, arrowheads), whereas the segmental expression domains were never recovered.

**Figure 3.**
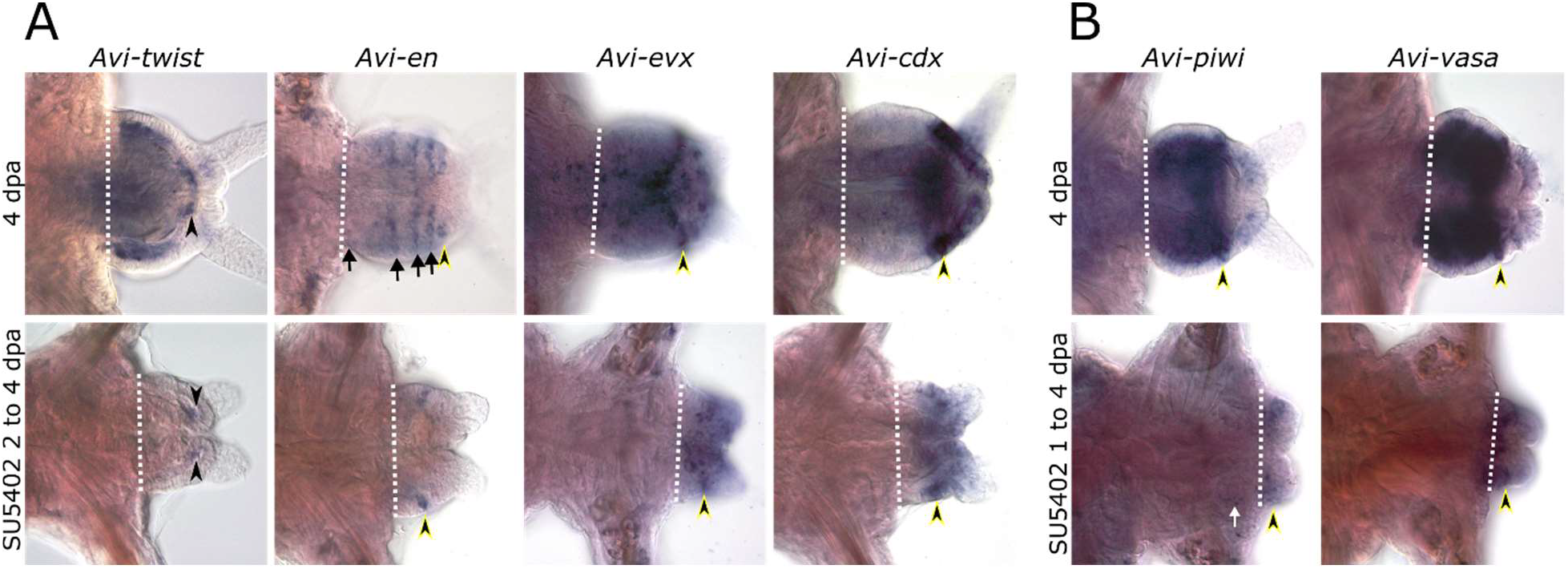
Posterior regeneration at 4 dpa stage in DMSO controls (upper row) and after inhibition of FGF pathway (lower row) starting at 2 dpa (A) or 1 dpa stage (B). All planes are ventral views. Dotted line indicates amputation plane, arrowheads with yellow outline point to the growth zone. In *Avi*-*twist* arrowheads indicate mesodermal domain in the circumpygidial muscles; in *Avi-en* arrows indicate expression stripes, where the one before the growth zone is discontinuous on the ventral side; in *Avi-evx* arrow points to signal in the nervous system (VNC) and arrowhead points to the expression in the posterior growth zone; in *Avi-piwi* white arrow points to the signal on the level of muscles in the last old segment.

*Avi-en* expression at 4 dpa manifests as metameric transverse stripes in superficial epidermis and underlying mesenteries. The anterior-most dashed stripe demarcates the boundary between the last old segment and the regenerate, two middle stripes are continuous at the ventral and lateral sides of the regenerate (Fig. 3A, arrows), and two posterior stripes disconnect at the ventral midline (Fig. 3A, arrowhead with yellow outline indicates the posterior growth zone). After two days of FGFR inhibition at 2–4 dpa we reliably detected only one discontinuous stripe of *Avi-en* expression, which presumably corresponds to the posterior-most domain in the growth zone (Fig. 3A, arrowhead). This domain is characterized by much weaker staining (compared with DMSO controls) localized in both ectodermal and mesodermal layers at the ventrolateral sides just in front of the pygidial circular blood sinus. Occasionally, at the border between the old segment and the regenerate, we observed individual spots of *Avi-en* expression at the limit of the method’s resolution. This anterior-most transverse domain is normally the only one established by 2 dpa, when we started inhibition. Thus, our experimental treatment suppressed the expression of the anterior *Avi-en* stripe but did not prevent the emergence of the posterior stripe presumably corresponding to the growth zone.

Aiming to clarify the identity of the posterior territories of the regenerative bud we used *Avi-evx*, which normally operates in the growth zone (specifically in the ring of superficial cells), pygidium and hindgut, as well as in some ventral neural cells of the forming segments (Fig. 3A, arrow). After using SU5402 at 2–4 dpa this differential expression pattern became less apparent. The brightest signal was observed in a slightly disorganized transverse stripe of superficial cells of the growth zone (Fig. 3A, arrowhead). Behind this stripe, we observed less intense staining in the pygidium and hindgut. A few scattered *Avi-evx*-positive cells on the anterior ventromedial part of the regenerate most likely corresponded to neural precursors. Similar to *Avi-evx*, we did not observe significant differences between control and experimental patterns of *Avi-cdx* expression. Despite the disproportion of the regenerate, experimental specimens demonstrated relatively undisturbed *Avi-cdx* expression in the whole posterior half, including the growth zone (distinguishable by more intense staining), pygidial epithelium and hindgut (Fig. 3A). Altogether these data indicate that the anterior part of the regenerative bud, which gives rise to segments, is severely affected by FGFR suppression, so it becomes unable to establish identity of mesodermal and ectodermal tissues. In contrast, the posterior regenerate’s part, which forms by 2 dpa stage and contains the growth zone and pygidium, is mostly unaffected after inhibiting FGFR at 2–4 dpa.

Since we detected the expression of positional markers of the growth zone, inhibiting FGFR at 2– 4 dpa (Fig. 3A), and the dedifferentiation signatures of its progenitor cells were not impaired inhibiting FGFR after the first dpa (Fig. 2, lower row), we decided to check whether the growth zone is formed under inhibition at 1–4 dpa. In other words, whether the multipotent state obtained by some cells at 1 dpa stage remains after pharmacological inhibition, so those cells may give rise to the multipotent cell fates of the growth zone. We chose to test *Avi-vasa* and *Avi-piwi* as candidate genes for this experiment, because of their predominant expression in the growth zone as markers of multipotency. At 4 dpa stage both genes have almost identical expression patterns, including the ectodermal and mesodermal parts of the growth zone, mesodermal segmental tissues, and pygidial ectoderm (Fig. 3B). Following FGFR inhibition at 1–4 dpa we observed an extremely small and hypomorphic regenerative bud, resembling that of 2 dpa stage. Expression of *Avi-vasa* and *Avi-piwi* in experimental specimens was disturbed, making these two patterns significantly different from each other. A common domain was confined to slightly dispersed superficial and internal cells localized to circular zone in the middle part of the bud, which we interpreted as a restored posterior growth zone. *Avi-piwi*-positive cells were additionally found in the adjacent old segment at the focal level of muscle cells (Fig. 3B, arrow), resembling those after inhibition at 1–2 dpa (Fig. 2, lower row). *Avi-vasa* demonstrated wider expression in the regenerate, including both ectodermal and mesodermal layers (Fig. 3B). In contrast to DMSO controls, the mesodermal tissues of experimental specimens had mosaic *Avi-vasa* signal. Thus, transformation of

*Avi-vasa* and *Avi-piwi* expression patterns (achieved by 1 dpa in normal conditions) during the next 3 days of inhibition indicated the preservation of the multipotent status of cells within the boundaries of the restored growth zone.

## Discussion

Our study examined different stages of regeneration, allowing us to identify key FGF-dependent processes (Fig. 4). We provide new evidence that in *A. virens* regeneration, FGF signaling is involved in blastema induction by stimulating prior dedifferentiation, proliferation of its cells, control of differentiation of mesodermal derivatives in the anterior regenerate, and the formation of segment boundaries. Some of these functions of FGF signaling are found in representatives of different taxa, indicating evolutionary conservatism of certain roles of this signaling pathway, such as induction of the mesodermal cells and their derivatives. Other functions, in particular those associated with posteriorization, on the contrary, seem to be an evolutionary acquisition of vertebrate FGFs.

**Figure 4.**
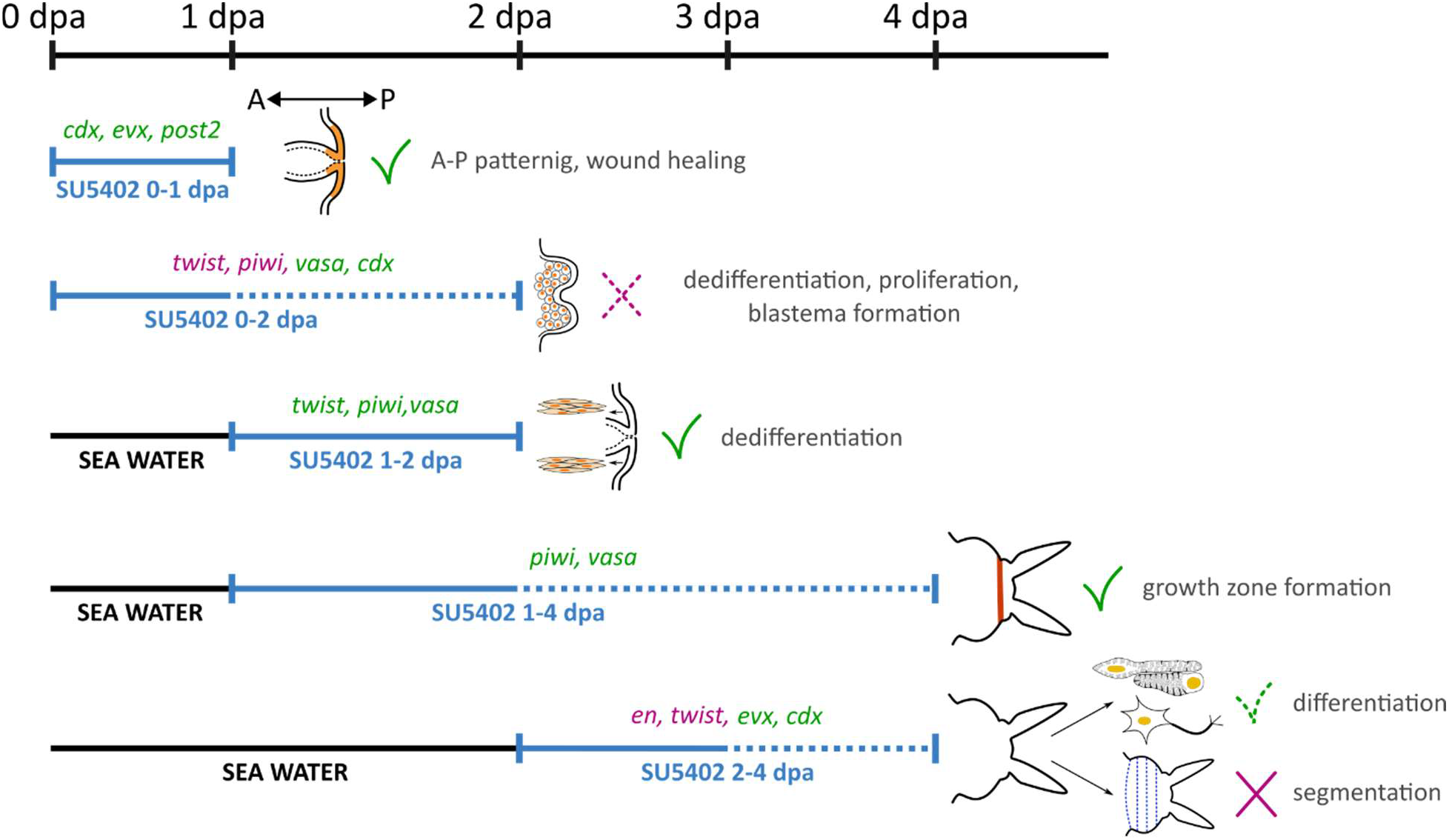
Effects of FGFR suppression on key processes during *A. virens* regeneration from amputation (0 dpa) up to 4 dpa. The period of FGFR inhibition is indicated by the blue sector, while prior cultivation in seawater is represented by the black line. The dotted portion of the blue sector indicates the time period during which FGFR inhibition results in a delay in the rate of regeneration. FGF-dependent processes include local cell dedifferentiation, proliferation, blastema formation, and segmentation of the regenerative bud (marked by magenta X; a dotted line indicates a partial effect). These processes were identified through gene expression dynamics, with the corresponding genes listed in magenta above the blue sector. Other events, such as anterior-posterior patterning, multipotency maintenance after the formation of the wound epithelium, posterior growth zone formation, and tissue differentiation, are FGF-independent (indicated by green check marks). However, differentiation of segmental components of the musculature and nervous system requires FGF activity, while pygidial patterning does not (indicated by a dotted check mark).

Comparing the role of FGF signaling in embryonic development with reparative processes can provide valuable insights into the evolutionary origins of regeneration mechanisms (Czarkwiani et al., 2021). Investigation of the conserved and divergent functions of this signaling pathway across different taxa can shed light on the shared developmental strategies that underlie both embryonic patterning and the re-establishment of lost structures during regeneration. *A. virens* and related polychaetes provide a valuable model system for studying this issue (Fischer, 1999; Özpolat et al., 2021). This comparative approach may elucidate the fundamental principles governing regenerative capacities and reveal the evolutionary adaptations that have enabled some species to maintain robust regenerative abilities, while others have lost this capacity.

### Blastema and mesoderm induction rely on FGF signaling

Our experiments revealed the requirement of FGF signaling for blastema induction at the earliest time period of restorative process. This is demonstrated by the altered expression patterns of the multipotency markers *piwi* and *vasa* following FGFR inhibition. Blastema formation is a crucial part of successful regeneration in nereidid worms, and the blastemal cells originate from the dedifferentiated tissues of the wound-adjacent segment (Boilly, 1969; Hill, 1970; Kozin & Kostyuchenko, 2015; Planques et al., 2019; Kostyuchenko & Kozin, 2021; Paré et al., 2023; Stockinger et al., 2024; Bideau et al., 2024). Experiments with EdU labeling in *A. virens* (Shalaeva & Kozin, 2023) and *Platynereis dumerilii* (Bideau et al., 2024; Planques et al., 2019) provide direct evidence that the blastema consists of cells activated in response to amputation within the wound-adjacent segment. Molecular data, such as the de novo expression of GMP genes post-amputation in *A. virens*, further supports the dedifferentiation hypothesis. Our previous work showed that the FGF pathway plays a role in blastema formation via proliferation control, with FGFRs expressed in the wound epithelium and later in the blastemal cells. Similar to *A. virens*, a potential competence for FGF signaling has been noted in the cellular sources of regeneration of planarians and vertebrates (Agata & Umesono, 2008; Maddaluno et al., 2017). Transcriptomic data have revealed overexpression of FGF components during regeneration and fission in several annelids (Bhambri et al., 2018; del Olmo et al., 2022; Nyberg et al., 2012; Paré et al., 2023; Ribeiro et al., 2019). Single-cell RNA-seq analysis of asexually reproducing oligochaete *Pristina leidyi* further supports an FGF-mediated regulation of proliferative precursors of new tissues, since the FGFR transcript was significantly enriched in *piwi*+ cells (Álvarez-Campos et al., 2024).

In the current study, we used molecular markers expressed in the blastema and wound epithelium to address whether these regulatory genes are activated by the FGF pathway. The absence of *piwi* and *twist* expression after FGFR inhibition immediately after amputation until the blastema formation stage (0–2 dpa), indicates the FGF-dependency of these processes (Fig. 2). In contrast, the induction of the multipotency marker *vasa* in the wound and adjacent segment suggests that the functions of *vasa* and *piwi* may differ in the pool of dedifferentiating cells in *A. virens*, a topic that warrants further investigation. Further research into the expression of *nanos*, another key player in the germline multipotency program within undifferentiated cells of *A. virens* (Kostyuchenko, 2022), could shed light on the processes of dedifferentiation, proliferation, and subsequent differentiation occurring at the wound site under the FGF control. Examining susceptibility of *pumilio* and *myc* expression, reported to be involved in regeneration or acting alongside *nanos* (Planques et al., 2019; Kostyuchenko et al., 2023), could provide further insights into these processes. Anyway, the revealed expression of *piwi*, which is sensitive to the timing of FGFR inhibition, supports a role for this gene in proliferation control, as corroborated by our previous EdU labeling experiments (Shalaeva et al., 2021; Shalaeva & Kozin, 2023). Overall, the presented findings highlight the crucial requirement of FGF signaling for the induction of the cellular sources that drive regeneration in *A. virens*. The published transcriptomic data (Álvarez-Campos et al., 2024; Bhambri et al., 2018; Paré et al., 2023; Ribeiro et al., 2019) indicate that this interpretation is likely to be valid for other annelid species.

Mesoderm specification is a key process regulated by the FGF pathway during deuterostomes’ development (Amaya et al., 1991; Kimelman & Kirschner, 1987; McFann et al., 2022), and inhibition of FGFR blocks mesoderm formation (Amaya et al., 1993; Fan et al., 2018; Green et al., 2013; Kim & Nishida, 2001; Schulte-Merker & Smith, 1995). A similar FGF-dependent mechanism has been observed in regeneration (see Introduction). In zebrafish fin regeneration, FGFR suppression immediately after amputation leads to wound closure without blastema induction and proliferation (Poss et al., 2000), similar to the findings in *A. virens*. The mechanism of fin blastema induction involves the activation of Fgf20 in the wound epithelium, which stimulates the underlying mesenchyme (Shibata et al., 2016). The latter provides a local source of Fgf3 supporting further cellular divisions. In annelid regeneration, blastemal cells also originate from mesodermal precursors, and in *A. virens*, FGF ligands and receptors are transcribed in the wound epithelium and underlying blastemal precursors during the earliest stages (Shalaeva et al., 2021), suggesting possibility of a similar to zebrafish induction process. One of the well-known pan-bilaterian mesodermal markers is *twist*, and the FGF pathway is known to target *twist* (Fan et al., 2018; Imai et al., 2003; Zhao et al., 2020) as well as other bHLH transcription factors (Fletcher & Harland, 2008; Row et al., 2018). Our study demonstrated the FGF requirement for the activation of *twist* expression during the first days after amputation.

Furthermore, the induction and patterning of mesoderm via the FGF pathway is suggested to be ancestral in the development of several spiralian taxa (Andrikou & Hejnol, 2021; Seudre et al., 2022; Tan et al., 2023). In the phoronid *Phoronopsis harmeri* embryos, whose *fgfr* and *twist* are coexpressed in mesoderm, suppression of FGFR prevents *twist* activation and result in visible mesoderm malformations (Andrikou & Hejnol, 2021). Similarly, annelid development relies on FGF signaling for mesoderm induction, particularly through the MAPK/ERK downstream pathway. In *A. virens* development, metameric mesoderm formation was disrupted using the pharmacological agent U0126, which suppresses ERK1/2 (Kozin et al., 2016). In the basal annelid *Owenia fusiformis*, the FGF pathway induces mesoderm formation, activating downstream genes such as *twist*. Its expression domain disappears from *Owenia* mesoderm after inhibition of either FGFR or ERK1/2 (Seudre et al., 2022).

Our results indicate that FGF signaling is crucial not only for inducing the mesoderm (Fig. 2, 3A, 4), but also for specifying some of its parts and muscle derivatives within the zone where new segments form (Shalaeva et al., 2021). This process mirrors embryonic and larval development, where the mesoderm depends on FGF signaling and may utilize similar downstream genes. The loss of the segmental *twist* domain, paired with the maintenance of the most terminal one after FGFR inhibition at 1–4 dpa, further supports the suggested heterogeneity within the early regenerative bud of *A. virens* (Shalaeva & Kozin, 2023). Specifically, the most posterior part of the mesoderm presumably segregates very early and provides material for the circumpygidial muscle ring independent of the FGF pathway. Collectively these data propose that the FGF-dependent stimulation of mesodermal tissues during regeneration can be explained as a redeployment of the evolutionarily conserved embryonic gene circuitry.

### Anterior parts of the regenerative bud are FGF-dependent, posterior parts are not

The role of FGF signaling in maintaining the AP axis is well-established in vertebrate development and regeneration, with posteriorization being a specific function in the trunk and tail regions (Christen & Slack, 1997; Dorey & Amaya, 2010; Griffin et al., 1995). However, in the developing neural tube of lower chordates like ascidians, FGF signaling acts at the anterior body end (Wagner & Levine, 2012). In our study on *A. virens* regeneration, which involves repatterning of the AP axis (Novikova et al., 2013), we did not find evidence to support a role for FGFs in posteriorization. Instead, we observed that posterior genes such as *cdx, post2*, and *evx* were able to establish their expression domains even when FGF signaling was pharmacologically inhibited (Fig. 1). This suggests that neither the initial wound healing nor the re-establishment of the posterior-most identities, which are part of the morphallactic changes, are dependent on FGF signaling in *A. virens*. On the contrary, the revealed connection between FGF signaling and epimorphic processes, potentially mediated by regulation of cell proliferation, multipotency, adhesion, and metabolism (Maddaluno et al., 2017; Mossahebi-Mohammadi et al., 2020; Oginuma et al., 2017; Ornitz & Itoh, 2022), suggests that the evolution of FGF involvement in the posterior axial elongation of vertebrates may be explained by the expansion of the zone of multipotent precursors at the posterior end of the embryo.

Our experiments revealed that regenerating areas located along the AP axis have different dependences on FGF signaling. The regenerative bud of nereid annelids comprises three distinct regions: the most posterior part, the pygidium; the posterior growth zone (GZ) in front of the pygidium, which provides cellular material for segment restoration; and the most anterior part, where new segments mature (Kozin et al., 2017; Kostyuchenko & Kozin, 2021; Planques et al., 2019). The segmental part of the bud and GZ demonstrate higher levels of cellular proliferation (Shalaeva & Kozin, 2023) and transcriptional activity (Niwa et al., 2013; Stockinger et al., 2024), including expression of multipotency markers (Kozin & Kostyuchenko, 2015). Our previous findings in *A. virens* showed that by 4 dpa, *Avi-fgfr1* is expressed in the mesoderm of the bud, while *Avi-fgfr2* has ectodermal expression, and both FGF ligands, *Avi-fgf8/17/18* and *Avi-fgfA*, have dual specificity in this regard (Shalaeva et al., 2021). In contrast, the pygidium is fully formed by this time and largely quiescent in terms of proliferation and gene expression. Inhibiting FGFR after blastema formation (at 2–4 dpa) disturbs the morphogenesis of pygidial cirri, but the overall organization of the pygidial nerves and muscles (Shalaeva et al., 2021), as well as the expression of posterior genes (*evx, cdx*), remains largely unchanged (Fig. 3). In line with that, by 3–5 dpa *Avi-fgfr1* and *Avi-fgfr2* expression clears from pygidial mesodermal and ectodermal tissues, respectively (Shalaeva et al., 2021). Taken together with the recovered expression of *piwi, vasa, cdx*, and *en* in the GZ, it indicates the ability to achieve molecular identity of both the pygidium and GZ, even when FGFR inhibition is initiated at 1 dpa (Fig. 3).

The discussed data suggest that the establishment of posterior positional information during *A. virens* regeneration occurs independently of FGF signaling, unlike in vertebrate mesodermal and neural tissues (Aulehla & Pourquié, 2010; Villanueva et al., 2002). In contrast, the FGF pathway appears to be most crucial for the segment formation area. While the absence of *twist* and *en* expression domains in this area could be attributed to a lack of regenerated tissues, our previous experiments with SU5402 exposure at 2–4 dpa demonstrated uniform EdU labeling throughout the regenerative bud (Shalaeva et al., 2021). This indicates that cell proliferation doesn’t stop completely, providing certainly reduced cellular material. Therefore, some aspects of axial elongation appear to be beyond the FGF control in our model. However, the tissue generated anterior to the pygidium fails to undergo commitment and differentiation upon FGFR inhibition.

According to our results, FGF signaling appears to play a critical role in the development of the anterior part of the regenerative bud, particularly in the formation of segmental boundaries and the differentiation of tissues within the segmental region. This is clearly illustrated by the loss of *en* expression, a marker of segmental borders, and the absence of metameric *twist* expression in the mesoderm. This aligns with findings from other regeneration models, such as *Hydra* (Lange et al., 2014; Sudhop et al., 2004), where FGF signaling contributes to boundary formation. Moreover, this function is also seen in the vertebrate neural tube, where FGF regulates *en* expression at the midbrain-hindbrain boundary (Gibbs et al., 2017; Scholpp et al., 2003). Similar to these models, we observed that FGFR inhibition disrupted the anterior-most *en* stripe at the boundary between the old and regenerating tissue, suggesting that FGF signaling is crucial for maintaining the anterior domain of the regenerative bud. This role of FGF is consistent with its action in other invertebrates, where FGF signaling is often localized to anterior structures during regeneration. For example, in planarians, FGF signaling is crucial for anterior brain regeneration (Cebrià et al., 2002; Ogawa et al., 2002), and in *Dugesia japonica, Djfgf* antagonizes Wnts to establish the AP axis (Auwal et al., 2020). Similarly, in *Amphioxus*, FGF signaling regulates the formation of the most anterior somites (Bertrand et al., 2011). These observations imply that FGF signaling may play a shared role in regulating anterior patterning during regeneration and embryonic development of metazoans, although further research is required to confirm this potential evolutionary conservation.

In summary, our results highlight the critical role of FGF signaling in maintaining mesodermal identity and facilitating metamerization within the regenerative bud of *A. virens*. However, neither the initial wound healing nor the re-establishment of posterior-most cell fates depends on FGF activity. Its area of responsibility is limited to the anterior part of the regenerative bud, where FGFs play a crucial role in segmental boundary formation and tissue differentiation. These features of FGF signaling appear to be evolutionary conserved across metazoans, as similar roles have been observed in development and regeneration of other species.

## Acknowledgements

This research was funded by the RSF grant 23-74-10046, https://rscf.ru/en/project/23-74-10046/. We are grateful to the research resource centers “Microscopy and Microanalysis”, “Molecular and Cell Technologies”, “Chromas”, “Culture Collection of Microorganisms”, and the Marine Biological Station of St. Petersburg State University for their technical support.

## Notes

### Competing Interest Statement

The authors have declared no competing interest.

